# HOPE: Interpretable Histology Analysis with Spatial Omics-Derived Signatures for Precision Oncology

**DOI:** 10.64898/2026.06.03.729847

**Authors:** Tianyi Wang, Matthew Bieniosek, Tamara J. Krpicak, Mingyuan Luan, Benjamin Ruf, Christian M. Schürch, Aaron T. Mayer, Ruibang Luo, Alexandro E. Trevino, Zhenqin Wu

**Affiliations:** School of Computing and Data Science, The University of Hong Kong, Pok Fu Lam, Hong Kong SAR; Enable Medicine, Menlo Park 94025, CA USA; Department of Bioengineering, University of Pittsburgh, Pittsburgh 15260, PA USA; Department of Internal Medicine I, University Hospital Tübingen, Eberhard Karls University of Tübingen, Tübingen, Germany; M3 Research Center for Malignome, Metabolome and Microbiome, Faculty of Medicine, University Tübingen, Tübingen, Germany; Department of Pathology and Neuropathology, University Hospital and Comprehensive Cancer Center Tübingen, Germany; Cluster of Excellence iFIT (EXC 2180) “Image-Guided and Functionally Instructed Tumor Therapies”, University of Tübingen, Germany

## Abstract

Hematoxylin and eosin (H&E) stained images are fundamental clinical tools for disease assessment. However, even with advanced computational models, their prognostic capabilities remain limited. Spatial omics characterizes tumor microenvironments (TME) in detail yet remains clinically inaccessible due to cost and complexity. In this study, we present HOPE, a lightweight framework that learns TME signatures from paired H&E and spatial omics data during training, then applies these to H&E alone at inference. Leveraging H&E foundation models, HOPE consistently outperforms identical architectures trained without spatial omics guidance across cancer types and cohorts. It further generates interpretable annotations of TME signature on H&E regions, stratifying patients into biologically coherent groups with different prognostic outcomes. HOPE establishes a practical route to translate high-content spatial omics discoveries into scalable, clinically deployable tools.

## 1. Introduction

Accurate clinical characterization improves patient outcomes and may help inform personalized treatment decisions. The analysis of Hematoxylin and Eosin (H&E)-stained images is a widespread, effective approach to diagnosing and classifying disease based on microscopic tissue structures [1]. Recently, taking inspiration from rapid advances in computer vision, computational methods have emerged to automate these pathology workflows. Artificial intelligence (AI)-powered foundation models, trained on massive histology datasets [2, 3] or image-text pairs [4, 5], have achieved impressive performance in clinical tasks [6] like cancer diagnosis, subtyping, prognosis and response to therapy prediction, survival modeling, and more. However, H&E images alone lack the molecular information necessary to fully characterize tumor microenvironments (TMEs), limiting the scope of these tools as biomarkers for next generation therapeutic strategies.

Spatial omics technologies, including spatial transcriptomics [7, 8] and proteomics [9, 10], enable richer *in-situ* measurements like molecular marker expression and context-aware analysis of cells within native tissue environments [11]. These technologies have been applied to reveal molecular patterns in tissue and their clinical relevance to cancer progression or treatment outcomes [12–15]. AI models trained on spatial omics inputs have also been used for clinical tasks such as predicting patient prognosis and treatment response [16–20]. These approaches have yielded more nuanced molecular signatures associated with clinical outcomes. However, the high cost of spatial omics creates a barrier to applying these biomarkers in clinical settings.

Recognizing this limitation, recent research has moved towards joint analysis of spatial omics and H&E images. Early representative works enhanced spatial domain detection by integrating H&E-derived tissue morphology [21]. Later studies demonstrated the enhanced predictive power of direct multi-modal modeling across H&E and diverse spatial omics modalities [22, 23]. Nevertheless, these tools still require spatial omics input during inference, limiting their feasibility for routine clinical application.

Other approaches instead try to infer more complex biomarkers, including gene expression [24–26], protein expression [27, 28], and coarse-grained cell types [29–31], directly from H&E images. Although promising, whether they can generalize across staining or institutional batch effects remains uncertain. More importantly, the biomarkers with the strongest inference results are often too superficial for nuanced disease assessment. Recent spatial omics studies emphasize this caveat [32, 33], showing how disease or treatment courses are often driven by complex, multi-scale interactions within the TME, rather than isolated features.

To address the gap between powerful prognosis-associated signatures revealed from costly spatial omics and clinically ubiquitous H&E images, we propose HOPE (HistOlogy model with spatial Proteomics Enrichment), a framework that transfers omics-derived TME insights to H&E and identifies those insights from H&E alone at inference time. To demonstrate our method, we leverage prognosis-associated TME labels distilled from spatial omics studies to supervise H&E-based predictors in a multitask setting, enabling both improved prognosis predictions and biologically grounded interpretability. Evaluated across diverse, multi-institutional patient cohorts, HOPE achieves consistent performance improvements in both survival and treatment response predictions, identifies coherent TME patterns linked to prognosis, and maintains robust generalizability across settings.

## 2. Results

### 2.1. Overview of HOPE

HOPE is a lightweight framework designed to transfer disease-associated signatures—broadly defined as spatially resolved molecular or cellular patterns linked to clinical characteristics—discovered from spatial omics analysis to H&E images (Fig. 1). The HOPE workflow consists of two phases: training and inference.

**Fig. 1.**
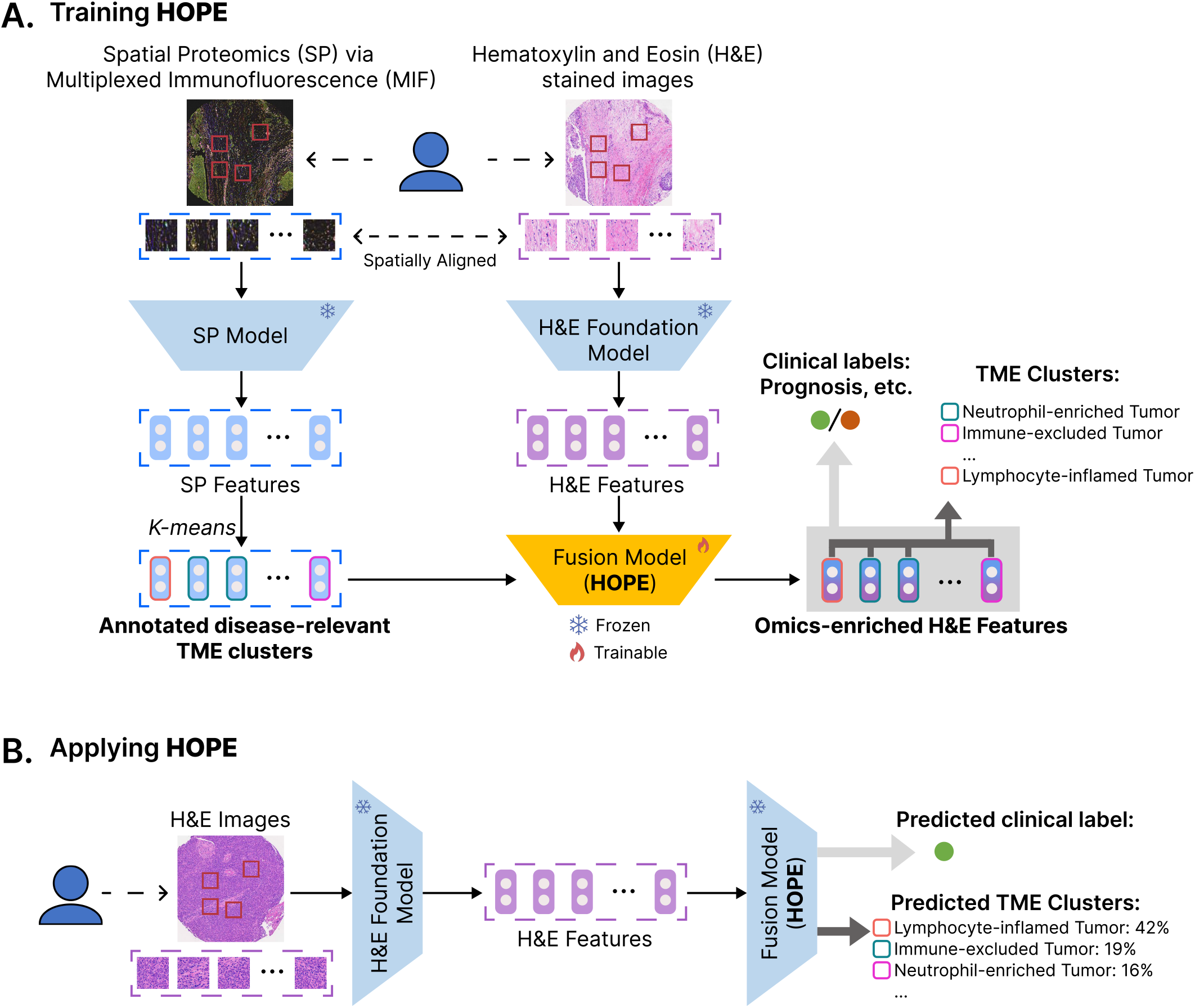
Overview of HOPE. **A**. Training workflow. HOPE integrates aligned spatial omics modalities with H&E-stained sections. Both modalities are partitioned into patches or units of equivalent size. Patch features and key disease-relevant TME clusters are extracted from the spatial omics modality using a specialized model. These TME cluster annotations are assigned to the corresponding H&E patch images. H&E patches are processed through a pretrained pathology foundation model to extract H&E features. HOPE takes H&E features as input for simultaneous prediction of TME clusters and clinical labels. **B**. Inference workflow. During inference, only H&E images are required. Features are extracted using the same pathology foundation model as in training, and passed through the trained HOPE to directly predict patch-level TME clusters and patient-level clinical labels.

During the training phase, disease-associated signatures are first annotated in the spatial omics modalities and then mapped to the spatially-aligned H&E patches. HOPE is, in principle, compatible with any spatial omics data modalities and computational approaches capable of extracting disease-associated signatures. Specifically in this study, we used multiplexed immunofluorescence (MIF) data alongside a dedicated AI approach [19] that captures the heterogeneity of cell neighborhoods in the TME. Leveraging this trained omics-specific AI model, key disease-relevant TME clusters, defined as groups of neighborhoods demonstrating strong associations with patient prognosis or treatment outcomes, are annotated and extracted. Such TME clusters are further mapped to the corresponding H&E image patches based on the spatially aligned H&E stain of MIF.

Next, HOPE establishes an H&E-only prediction pipeline to capture both the omics-derived disease-associated signature labels and the patient-level prognosis labels. H&E patch embeddings are first extracted using pathology foundation models. Throughout all experiments in this study, we used UNI2-h [2] by default unless specified otherwise. A multi-layer perceptron (MLP) head is then trained to jointly predict TME cluster labels and patient prognosis labels based on the H&E embeddings using a multi-task setting.

During the inference phase, HOPE operates solely on H&E images to perform inference of disease-relevant TME clusters and patient prognosis. In this phase, both the backbone pathology foundation model and the MLP adapter components are fixed. HOPE first generates patch-level predictions. Patient-level prognosis predictions are then derived via Multiple Instance Learning (MIL) aggregation. The final outputs consist of two elements: patient prognosis prediction and a spatially resolved map of TME clusters, which provides biological-interpretable anchors for the prognosis prediction.

### 2.2. Applying HOPE on head-and-neck cancer patients for prognostication

We demonstrated the HOPE workflow on two cohorts of head-and-neck cancer (HNC) patients. The goal is to infer patient prognosis after surgery treatment using only H&E stained images of the tumor sections, while identifying and locating key prognosis-associated signatures.

The HOPE pipeline was established on an HNC dataset of 81 patients collected at the University of Pittsburgh Medical Center (UPMC), containing spatially aligned H&E and MIF images stained from adjacent tissue sections (Methods). This dataset is hereafter referred to as UPMC-HNC. We first validated the HOPE workflow using a four-fold cross-validation strategy, during which patients were randomly divided into four groups. In each fold, both TME cluster discovery by the omics-specific AI model and the subsequent HOPE workflow were performed on three training groups, with validation conducted on the held-out group.

Spatial omics profiling of tumor tissue sections enabled cell annotation and extraction of disease-associated spatial signatures. As shown in Fig. 2A, the omics-specific AI model constructed cell graphs from MIF images and generated embeddings for subgraphs representing neighborhoods. These embeddings enabled annotations of key TME clusters that are strongly associated with prognosis, which were further paired with the corresponding H&E image patches.

**Fig. 2.**
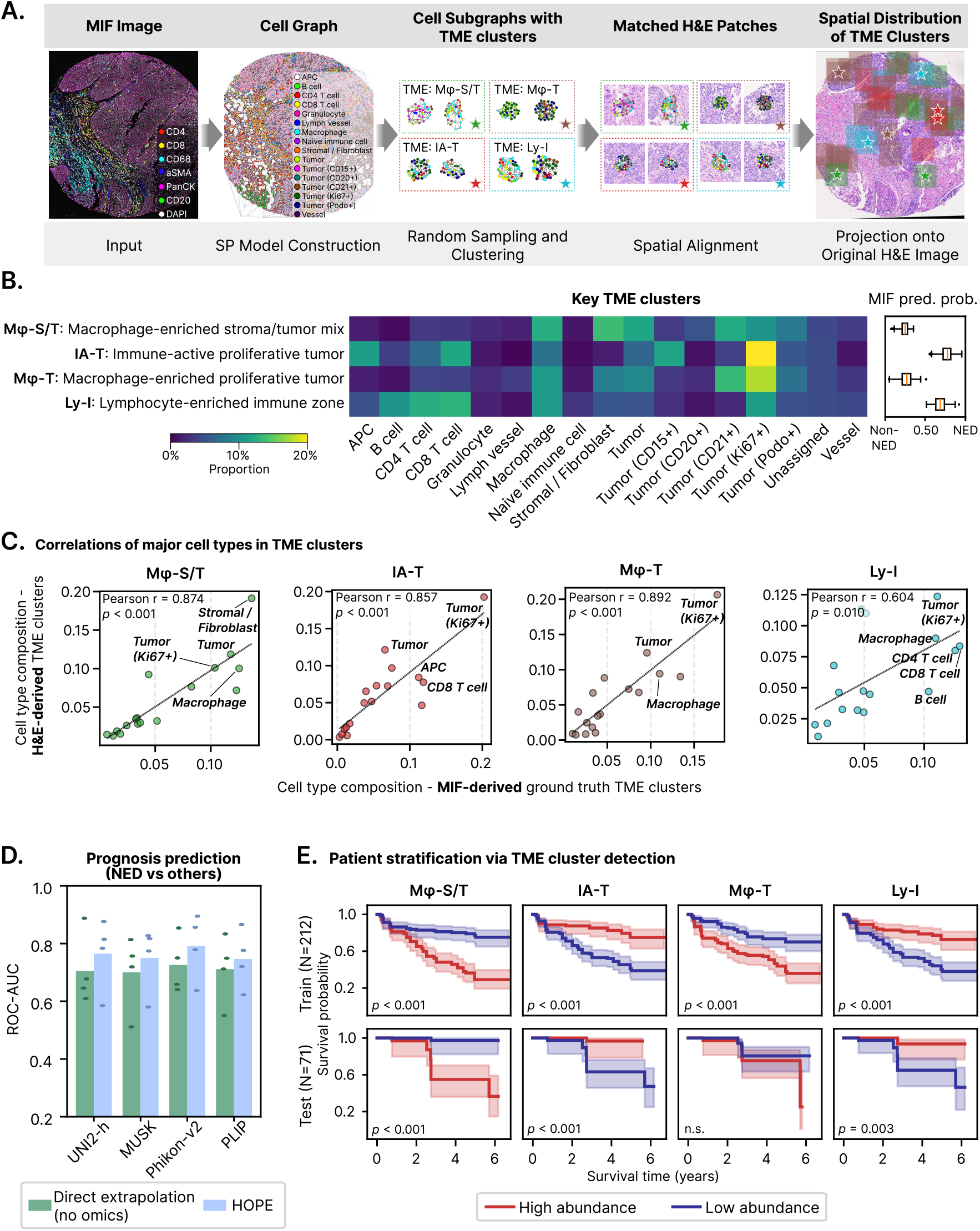
Survival prediction of UPMC-HNC dataset using HOPE. **A**. Spatial mapping of MIF-derived TME clusters on matched H&E patches. **B**. Cell type compositions of the TME clusters derived from the MIF-based AI model, with the right panel showing MIF-based cell graph prediction probabilities grouped by cluster for patient outcome prediction. **C**. Agreement between ground truth and HOPE-predicted clusters’ cell type compositions. **D**. Prognosis prediction performance (ROC-AUC) comparing HOPE with direct extrapolation. Patient-level predictions are aggregated from patch-level predictions by MIL. NED, no evidence of disease. **E**. Kaplan–Meier (KM) stratification by predicted TME cluster abundance, with individual H&E slides treated as independent samples to increase statistical power, showing associations between higher M*φ*-S/T or M*φ*-T and worse survival, and between higher IA-T or Ly-I and better survival.

In an example fold, four representative TME clusters—macrophage-enriched stroma/tumor mix (M*φ*-S/T), immune-active proliferative tumor (IA-T), macrophage-enriched proliferative tumor (M*φ*-T), and lymphocyte-enriched immune zone (Ly-I)—were identified as strong correlates of prognosis. Their cell compositions are summarized in Fig. 2B. Both macrophage-enriched clusters (M*φ*-S/T and M*φ*-T) are highly indicative of negative prognosis [34, 35]. IA-T mostly contains tumor tissues infiltrated by active T cells and antigen presenting cells, suggesting an active immune response to tumor and, consequently, better prognosis [36]. Ly-I contains local aggregates of B cells and T cells, pointing to the potential formation of tertiary lymphoid structures, also suggesting a positive prognosis [37].

The H&E-based HOPE workflow was subsequently established, which predicts patch-level TME clusters and patient-level prognosis for unseen test regions. To validate the TME cluster predictions, we assessed the agreements between cell type compositions of ground truth and predicted TME clusters in Fig. 2C. As TME cluster annotations were performed exclusively on training folds to prevent data leakage, ground truth labels were unavailable for test fold data. We evaluated prediction validity by computing Pearson correlations between cell type compositions of training and test folds. Training fold compositions were derived directly from ground truths (Fig. 2B). Test fold compositions were obtained on H&E patches assigned to the corresponding TME clusters by HOPE (Fig. S1). Across all four cases, we observed strong, consistent agreements, with Pearson correlation coefficients above 0.85 for M*φ*-S/T, IA-T, and M*φ*-T.

Inference outputs on two tumor cores are illustrated in Fig. 3. We leveraged the trained HOPE model to infer presence and abundance of key TME clusters in these two samples from patients with opposite prognosis: Example 1 comes from a patient with negative outcome (died of disease, DOD), whereas Example 2 comes from a patient with positive outcome (no evidence of disease, NED). These two cases exhibit distinct TME cluster distributions. Example 1 is characterized by the presence of M*φ*-S/T and M*φ*-T patches, with no detected IA-T or Ly-I. In contrast, Example 2 shows a reciprocal pattern where IA-T and Ly-I are abundant, while M*φ*-S/T and M*φ*-T are entirely absent. These results simultaneously lead to confident HOPE predictions at the patient level as negative and positive, respectively. Representative H&E patches from the four assigned TME clusters, which served as the vessels for transferring spatial omics-derived signatures to histology, are highlighted to demonstrate matched profiles between HOPE-generated H&E-based predictions and MIF-derived molecular ground truths.

**Fig. 3.**
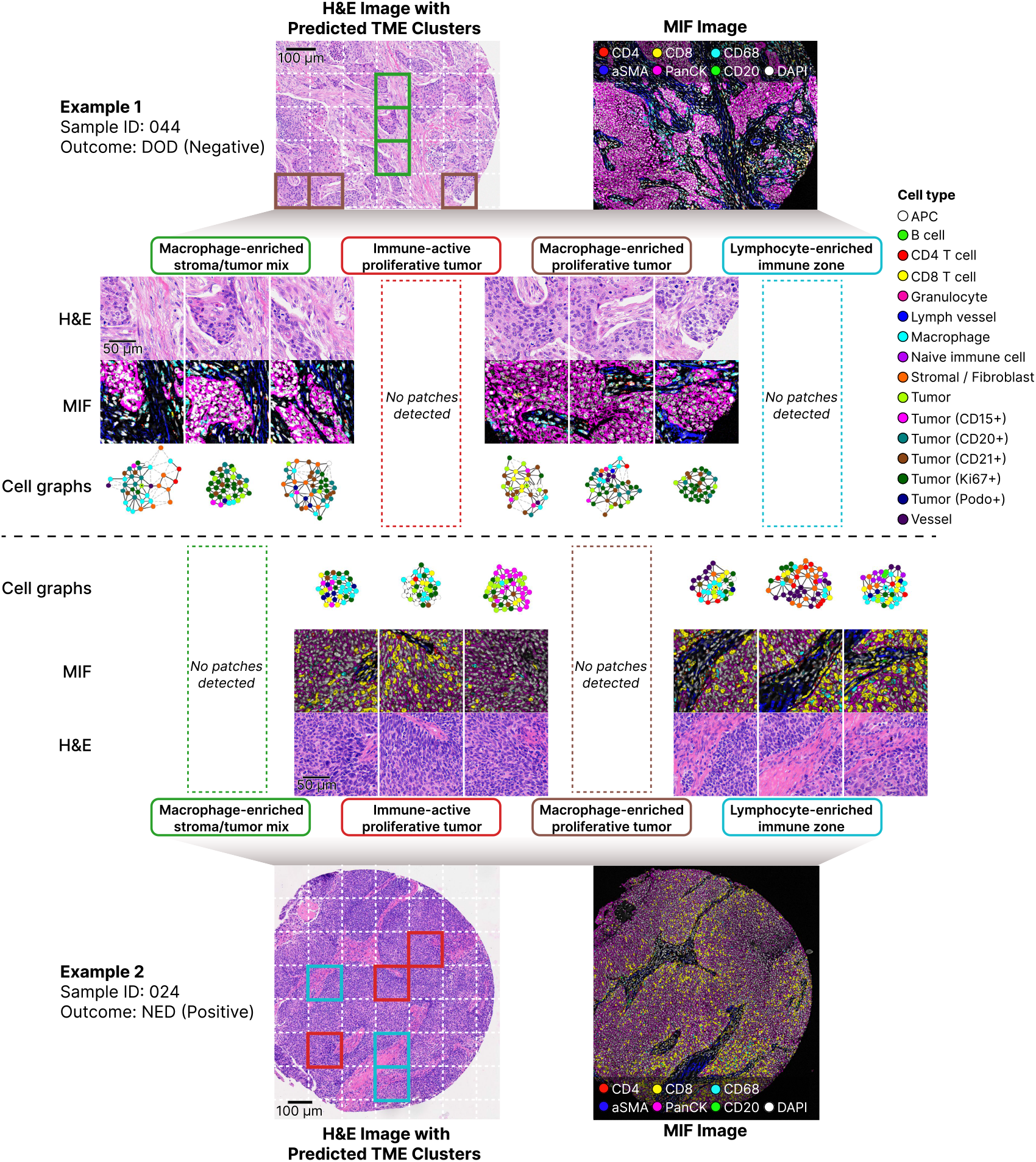
Representative cross-modality examples from UPMC-HNC. Two patients are shown to illustrate how MIF-derived spatial phenotypes are transferred to H&E. Example 1 is a negative outcome case (DOD), and Example 2 is a positive outcome case (NED). For each patient, matched cell graphs, MIF patches, and H&E patches are grouped by predicted TME cluster. Four representative clusters are visualized, and opposite spatial patterns are observed between the two examples.

Finally, we evaluated the overall performance of patient-level prognosis modeling (Fig. 2D) from HOPE. To assess robustness of HOPE, several pathology foundation models were employed as the backbone feature extractor, including UNI2-h [2], MUSK [4], Phikon-v2 [3], and PLIP [5]. Compared with direct extrapolation using solely patient prognosis labels, HOPE, with integration of the TME clusters, consistently improved ROC-AUC across all scenarios, with consistent performance gains of 3.5%–6.6% across diverse H&E encoders.

### 2.3. Biological interpretability from HOPE-predicted TME clusters

A key advantage of HOPE over direct extrapolation of pathology AI models is that HOPE conserves the interpretability of omics-derived signatures. Clinical decision-making prioritizes biologically interpretable features, while black-box AI models often cannot provide clear biological relevance for their outputs. HOPE addresses this challenge by augmenting predictions with disease-associated TME signatures, which come with clear molecular and cellular profiles.

We demonstrated this potential through patient stratification experiments (Fig. 2E). Using patch-wise annotations of TME clusters from HOPE, we bisected patients into low- and high-abundance groups for each TME cluster, with individual H&E slides treated as independent samples to increase statistical power. Higher abundances of M*φ*-S/T and M*φ*-T were significantly associated with lower overall survival (log-rank test, *p <* 0.001). Conversely, higher abundances of IA-T and Ly-I were significantly associated with better prognosis (log-rank test, *p <* 0.001). These trends held for test patients and were consistent across folds. Distributions of TME cluster abundances across patients with different prognosis labels showed similar results (Fig. S2): M*φ*-S/T and M*φ*-T proportions were markedly higher in non-NED patients, whereas IA-T and Ly-I were more abundant in NED patients.

### 2.4. Generalization validation for survival prediction

To further assess the generalizability of HOPE, we evaluated our model in a cross-institution application task. The tissue preparation, processing, staining, and imaging protocols for histology analysis vary across institutions, which often lead to poor generalizability of AI models. External validations on independent cohorts are therefore essential to distinguish predictive performance based on actual biological signals from dataset-specific biases.

To examine the added values of omics-derived insights for HOPE, we trained a HOPE pipeline using the aligned H&E and MIF data from the UPMC-HNC cohort, and evaluated it on HANCOCK [38], an independent, publicly available cohort of around three hundred HNC patients from the University Hospital Erlangen (Fig. 4A). Spatial signatures in the form of TME clusters were derived using the same pipeline described in Fig. 2A, employing the full UPMC-HNC cohort rather than only cross validation training folds. Representative TME clusters shown in Fig. 4B shared very similar compositions and indications as in our previous experiments. Using these TME clusters, HOPE was trained on UPMC-HNC and applied to HANCOCK H&E patches tiled from tumor centers.

**Fig. 4.**
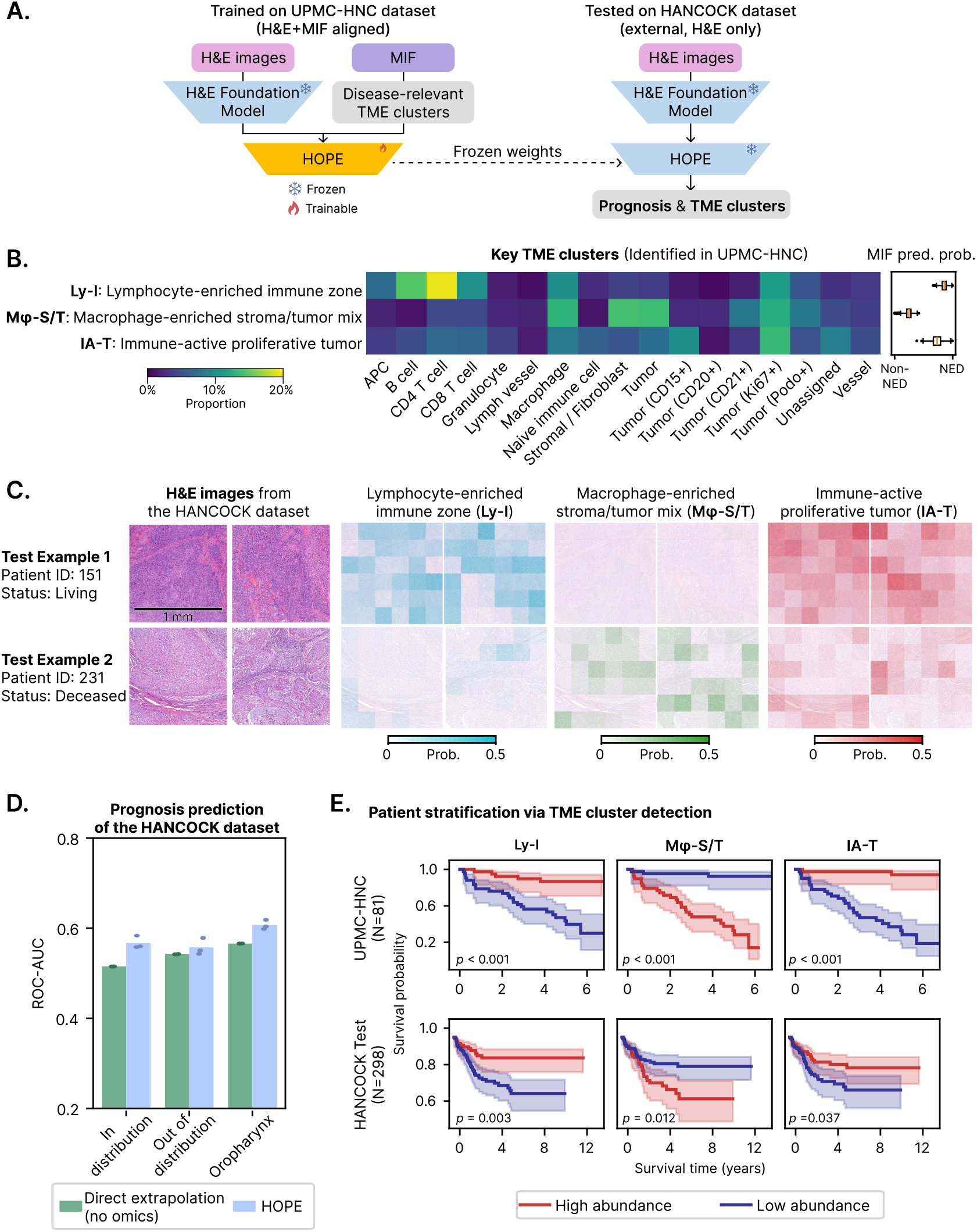
External generalization of HOPE on HANCOCK dataset for HNC prognosis prediction. **A**. Illustration of the cross-institution application task. HOPE is trained on UPMC-HNC and evaluated on an external dataset from a different institution. **B**. Cell type compositions of three key TME clusters derived using all UPMC-HNC data, with the right panel showing MIF-based cell graph prediction probabilities grouped by cluster for prognosis. **C**. Representative HANCOCK cases with patch-level predicted TME cluster probabilities, suggesting Ly-I/IA-T enrichment in a positive case and M*φ*-S/T enrichment in a negative case. **D**. ROC-AUC comparison between HOPE and direct extrapolation of pathology foundation model across three HANCOCK splits with three independent runs. **E**. Patient-level KM stratification by predicted TME cluster abundance in training and external test cohorts, where risk trends observed in UPMC-HNC were reproduced in HANCOCK.

We illustrate HOPE inference on the HANCOCK dataset with two representative H&E images, along with the predicted spatial maps of TME cluster probabilities (Fig. 4C). In an example region extracted from a patient with positive outcome (Example 1), immune-active clusters Ly-I and IA-T were more prominent. In the region (Example 2) from a deceased patient with negative outcome, more patches exhibited high M*φ*-S/T signals. Additional visual examples are illustrated in Fig. S3. For each TME cluster with well-defined cellular patterns derived from MIF analysis, the H&E patches identified in HANCOCK showed clear morphological similarity to the corresponding patches from UPMC-HNC. Quantitative prognosis modeling performances are presented in Fig. 4D, where we compared against direct extrapolation of pathology foundation model on three HANCOCK splits [38]. Across multiple independent runs, HOPE consistently achieved better performance in predicting survival. Notably, this is a challenging task due to patient distribution shift, differences in label definition, and treatment protocol heterogeneity between UPMC-HNC and HANCOCK.

We further demonstrated HOPE’s unique capabilities to infer key TME clusters through the stratification experiments in Fig. 4E. Results followed the same patterns: we noted positive associations between high Ly-I and IA-T abundance and survival, as well as negative associations between M*φ*-S/T abundance and survival. Highly significant separations are obtained in UPMC-HNC training data (log-rank test, *p <* 0.001) and remain reliable in the external HANCOCK cohort (log-rank test, *p <* 0.05). Distributions of TME cluster abundances (Fig. S4) supported the same findings. Survivors/NED patients exhibited higher Ly-I and IA-T proportions, while deceased/non-NED patients showed greater M*φ*-S/T abundance, with consistent observations in both UPMC-HNC and HANCOCK. These findings validated the generalizability of TME cluster-driven prognosis modeling approach of HOPE.

### 2.5. Applying HOPE to immunotherapy response prediction in hepatocellular carcinoma patients

Predicting treatment response is a critical goal in digital pathology. Accurate prognostic markers have the potential to directly guide effective therapeutic intervention and usher in more personalized care. Immunotherapy has become a transformative treatment option that improved survival outcomes for many malignancies [39], yet patient response rates remain highly variable [40], with limited reliable biomarkers [41]. To address this gap, we set out to validate the applicability of HOPE to predict patient response to immune checkpoint inhibitor (ICI) therapy in hepatocellular carcinoma (HCC), leveraging key disease-associated signatures identified from previous spatial multi-omics analysis.

We trained HOPE on a discovery HCC cohort with aligned H&E and spatial multi-omics data [20, 42] from a phase II clinical trial of ICI treatment for HCC. During the training phase, spatial signatures associated with immunotherapy response were directly derived from the source study of spatial multi-omics data, resulting in the annotations of seven major TME clusters: IA-S (immune-active stroma), Ly-T (lymphocyte-rich tumor), My-T (myeloid cell-rich tumor), Ec-T (endothelial cell-rich tumor), NST (non-specific tumor), LA (lymphoid aggregates), and InterFace (LA-tumor interface). As detailed in Fig. 5B, the cell type compositions of these clusters reveal distinct immune contexts across stroma-associated (IA-S, LA) and tumor-associated (Ly-T, My-T, Ec-T, NST) TME clusters. Notably, IA-S and Ly-T, both with abundant lymphocyte populations [43], exhibited high associations with immunotherapy response, whereas My-T and Ec-T, enriched with macrophages [44, 45] and endothelial cells [46], respectively, showed indications for resistance.

**Fig. 5.**
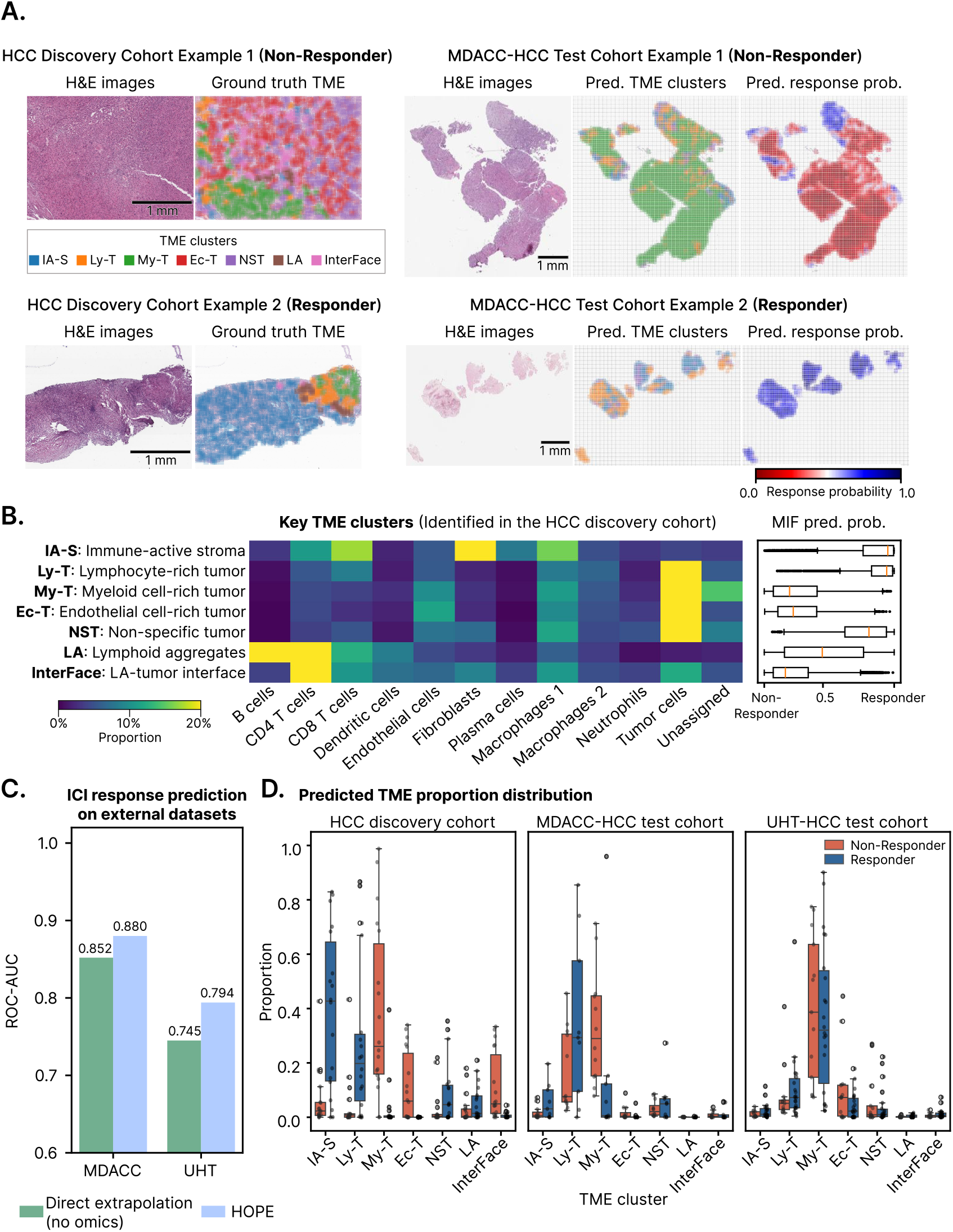
Immunotherapy response prediction with HOPE for hepatocellular carcinoma patients. **A**. Spatial distributions of TME clusters plotted for the discovery cohort samples (left) using ground truth, and for the external MDACC-HCC test samples (right) using HOPE predictions. The predicted maps of response probabilities for the test samples are also shown. My-T and Ec-T dominated non-responders, whereas IA-S and Ly-T dominated responders. **B**. Cell type compositions of TME clusters extracted from MIF data of the discovery cohort, with the right panel showing MIF-based cell graph prediction probabilities grouped by cluster for immunotherapy response. **C**. ROC-AUC comparison between HOPE and direct extrapolation of pathology foundation model across two external test cohorts on predicting immunotherapy response. **D**. Cohort-wise distributions of TME cluster abundances, summarized for seven major TME clusters.

For evaluation, we additionally obtained two external H&E-only cohorts from different institutions: MD Anderson Cancer Center (MDACC-HCC) and University Hospital Tübingen (UHT-HCC), consisting of 21 and 35 patient samples, respectively. While the recruitment criteria and treatment details for patients in these cohorts differed in some aspects (Methods), both datasets included response labels for pretreatment patient samples based on clinical assessments of therapeutic success.

We demonstrated qualitative prediction results in Fig. 5A. In the two examples from the discovery cohort (left), My-T and Ec-T dominated the non-responder tumor section, whereas IA-S dominated the responder section. When transferred to the external samples from MDACC-HCC (right), the same enrichment patterns were recovered: My-T dominated non-responder tumor sections, whereas IA-S and Ly-T dominated the responder section. Spatial maps of predicted immunotherapy response probabilities recapitulated the same spatial distributions as predicted TME clusters. More visual examples of inferred TME clusters from external H&E cohorts were illustrated in Fig. S5. Their consistency with corresponding H&E patches from the discovery cohort suggested that HOPE captures TME cluster-specific morphological patterns.

Quantitatively, HOPE achieved overall ROC-AUC performances of 0.880 and 0.794 on the two external H&E-only cohorts, showing improvements over direct extrapolation using the same pathology foundation model and training data (Fig. 5C). Fig. S6 shows additional results from an internal validation experiment, where we left out 30% of the spatial multi-omics patient samples from the discovery cohort for validation. Both TME cluster predictions and ICI response predictions (ROC-AUC of 0.946) yielded reliable performance.

HOPE further enabled predictions of specific response- and resistance-associated TME clusters in the external H&E-only data. Leveraging these outputs, we analyzed the patch-level abundances of these TME clusters. Consistent enrichment patterns were observed across the discovery cohort and both external cohorts. IA-S and Ly-T, both mapped to immune-active niches rich in lymphocytes, were more abundant in immunotherapy responders. Conversely, My-T and Ec-T, with notable immunosuppressive and angiogenic signatures in the spatial multi-omics analysis, were more abundant in non-responders (Fig. 5D). Among them, the predominant resistance-associated cluster of My-T further showed a stronger association with increased mortality (Fig. S7, log-rank test, *p* = 0.019) than manual assessment of response, which was largely determined by disease progression. Combined, these observations on biologically interpretable TME clusters justify the superior performance of HOPE and provide detailed, molecularly informed insights for H&E-only samples.

## 3. Discussion

To summarize, we presented HOPE, a lightweight framework that integrates spatial omics with histology to enable interpretable H&E-based prognostics. HOPE translates disease-associated signatures discovered from spatial omics into morphological patterns detectable in routine H&E images. This allows affordable, molecularly-informed biomarkers to be identified from limited spatially-aligned multimodal data and scalably deployed on larger cohorts with only H&E inputs. Crucially, HOPE obviates the need for expensive spatial omics measurements at test time—a nonstarter for practical clinical use—while still enabling the identification of key prognostic signals that have direct biological relevance.

Across multiple cancer types and prognostic tasks, HOPE consistently improved prediction performance over direct extrapolation with pathology foundation models, the state-of-the-art approach as of present. Perhaps more importantly, HOPE automatically provides biologically coherent interpretations for its predictions.

In the HNC case study, cross-validations on UPMC-HNC demonstrated that HOPE achieves consistent improvements over direct H&E extrapolation regardless of the choice of underlying pathology foundation model backbone. Prognosis-associated TME clusters, including higher-risk macrophage-rich clusters (M*φ*-S/T and M*φ*-T) and lower-risk immune-active clusters (IA-T, Ly-I), were identified and showed consistent patterns in both internal and external validation cohorts despite differences in patient demographics and treatment protocols.

In the HCC immunotherapy response modeling experiment, we showed that HOPE, trained with signatures annotated from spatial multi-omics data, consistently differentiated treatment responders from non-responders with only pretreatment H&E inputs. HOPE simultaneously identified the TME signatures most likely to associate with response or non-response, such as the contrasting lymphocyte-rich (Ly-T) and myeloid cell-rich tumor microenvironments (My-T) found in both discovery and external validation cohorts.

HOPE is a practical path for leveraging any spatial omics-derived biological annotations in routine digital pathology analysis. In other words, any biologically-meaningful derivatives of spatial omics analysis—gene expression programs, cell-cell interactions, spatial neighborhoods, and more—can be used to define the inputs to HOPE and later predicted from routine histology images. The framework is compatible with all spatially-resolved profiling technologies so long as spatially-coregistered histology is available. The choice to decouple training-time omics-based supervision from inference-time H&E-only prediction—which we validate on external cohorts—demonstrates the potential for clinical scaling and biological interpretability of HOPE biomarkers.

Certain operational constraints still exist for HOPE. First, HOPE’s omics-based supervision relies on upstream spatial omics analyses; any bias in the curation and selection of TME signatures may propagate to the H&E model and affect its downstream utility. Second, despite validations that cut across institutional and clinical settings, case studies in this work focus on deployment within a single cancer type. Cross-cancer or pan-cancer transfer of TME signatures will need to address challenges stemming from the heterogeneity of tissues and disease conditions. Third, while HOPE localizes key TME signatures at the image patch level (around 100–150 *µm*), mechanisms to further pinpoint the contributing cells from H&E images would allow further interpretability. Pairing HOPE with models to infer gene expression or cell identity from H&E is one such possible mechanism.

Looking ahead, we envision multiple directions to expand HOPE’s utility. Strategies to integrate multiple discovery cohorts with diverse molecular readouts, such as spatial transcriptomics and genetic markers, will expand the breadth of disease-associated signatures compatible with HOPE training. Confidence-aware prediction with calibrated uncertainty estimates at the patch level would support more reliable interpretations. Finally, extending HOPE beyond H&E to additional routine pathology modalities, particularly immunohistochemical (IHC) staining, will broaden its applicability in real-world workflows. Ultimately, we anticipate HOPE serving as a versatile bridge that translates high-dimensional molecular discoveries into clinically deployable histology biomarkers.

## 4. Methods

### 4.1. Acquisition and preprocessing of UPMC-HNC dataset

UPMC-HNC contains 308 regions in tumor microarray core format (1 mm diameter) from 81 head-and-neck cancer patients (2 to 6 cores, median 3 cores, per patient), collected at University of Pittsburgh Medical Center. Phenocycler multiplexed imaging (0.3775 *µm*/pixel) with 43 biomarkers was performed by Enable Medicine, capturing 2.37 million cells (6,248 ± 3,889 cells per region). 283 regions from the 81 patients with corresponding H&E from adjacent sections and patient prognosis labels were used in this study for establishing the HOPE pipeline. We used the primary outcome label as the prognosis prediction target, where no evidence of disease (NED) was treated as the positive class, and all other outcomes (alive with disease, died of disease, etc.) were treated as the negative class.

### 4.2. Acquisition and preprocessing of HANCOCK dataset

The HANCOCK dataset is a comprehensive collection of head and neck squamous cell carcinoma samples acquired between 2005 and 2019 at University Hospital Erlangen, containing tissues that are formalin-fixed and paraffin-embedded (FFPE), followed by H&E-stained whole-slide sections scanning or tissue microarrays constructed from 1.5 mm diameter cylindrical cores sampled from the tumor center and invasive front (40x, around 0.26 *µm*/pixel).

Here, we used H&E images of tumor centers obtained from the pre-extracted tumor center cores. For each H&E image, the central 2400 ×2400 region was retained and tiled into 400× 400 patches using a sliding window approach without overlap. Survival status served as the prediction label, where patients with non-tumor-specific death were excluded. The positive class was defined as living and the negative class as deceased. After filtering out cases with blank or invalid H&E images, 298 patients remained for the evaluation.

### 4.3. Acquisition and preprocessing of HCC discovery dataset

The HCC discovery cohort was downloaded from a recent spatial multi-omics study on a phase II clinical trial that evaluated the safety of perioperative anti-PD1 and anti-CTLA4 treatments [20, 42]. Major pathological response (MPR), defined as greater than 70% tumor necrosis at 6 weeks, was used as the indicator for response and non-response. Samples from 13 patients, comprising 6 responders and 7 non-responders, were submitted for spatial multi-omics profiling and H&E staining. The dataset contained 48 images with 51-plex immunofluorescence and 22 samples with Visium spatial transcriptomics, out of which 36 MIF images from 11 patients have corresponding H&E staining from adjacent sections. We used the patient-level MPR-based response/non-response labels to establish the HOPE pipeline.

### 4.4 Acquisition and preprocessing of HCC validation dataset

Two independent HCC validation cohorts were sourced from different institutions to validate the HOPE pipeline: MDACC-HCC from MD Anderson Cancer Center contains 21 H&E samples from 13 patients receiving the same perioperative ICI treatment as patients in the discovery cohort. MPR was used to derive labels for response/non-response, resulting in 12 responder samples and 9 non-responder samples. UHT-HCC from University Hospital Tübingen contains 35 H&E samples from 33 patients with advanced, unresectable HCC, all collected prior to anti-PD-L1 and anti-VEGF treatment. Treatment response labels were manually curated based on clinical tumor progression, resulting in 22 responder samples and 13 non-responder samples.

For both cohorts, raw H&E images were preprocessed using an identical pipeline: for each H&E image, tissue-containing regions were first identified based on sufficient tissue presence. These valid regions were then tiled into patches via a sliding window approach. Empirically, patch sizes were set to 280 ×280 pixels for MDACC-HCC and 400× 400 pixels for UHT-HCC to adjust for image resolution, with a stride of half the patch size. Binary patient-level response and non-response labels were derived from both datasets, respectively. Overall survival data of the patients from UHT-HCC were used for the TME cluster stratification experiment.

### 4.5. Obtaining spatial omics-derived signatures

Spatial omics-derived signatures were obtained from MIF data aligned with H&E images. We employed SPACE-GM [19], a graph neural network (GNN)-based model, to extract cell subgraph embeddings from all cohorts. SPACE-GM constructs cell graphs with cells as nodes and cell types as features for each whole MIF region. These graphs are partitioned into multiple 3-hop subgraphs representing local cellular neighborhoods. Employing MIL on these subgraphs as inputs, we trained SPACE-GM for patient prognosis prediction to obtain embeddings for each subgraph.

For the UPMC-HNC cohort, we randomly sampled cell subgraphs from the training set and clustered these embeddings into *C*_aux_ clusters via K-means to derive prognosis-associated TME signatures, with *C*_aux_ as a hyperparameter. We assigned biological interpretations based on TME cell type composition analysis, identified representative subsets with strong prognostic associations, and mapped their labels to the corresponding H&E patch regions. Patch regions were centered at the subgraph’s center node with a fixed radius. Specifically, given that these cell subgraphs covered areas up to approximately 400 pixels, corresponding H&E patches were cropped to 400 ×400 pixels. To prevent data leakage during cross-validation, TME cluster annotations were derived exclusively from training set MIF subgraphs. Consequently, test set MIF data lack ground truth TME labels, precluding direct accuracy assessment; we instead evaluated prediction validity through correlations with cell type compositions.

For the HCC discovery cohort, we followed the per-cell neighborhood annotation scheme characterizing 10 major microenvironment classes from the original study [20]. We consolidated these into 7 functional categories for HOPE training: CD4 T cell-rich lymphoid aggregates and B cell-rich lymphoid aggregates were combined as the “LA” class; tumor interface and fibroblast interface were combined as the “InterFace” class; T/M*φ*-rich stroma and IFN-*γ* activated stroma were combined as the “IA-S” class. H&E patch regions corresponding to cells with these annotations were extracted following the same protocol as UPMC-HNC, with patch size set to 224 × 224 pixels to match the HCC discovery cohort’s imaging resolution.

### 4.6. Extraction of H&E image embeddings

We employed H&E foundation models to extract H&E embeddings, including UNI2-h [2], MUSK [4], Phikon-v2 [3], and PLIP [5]. For each model, we adopted the recommended patch size, image transformation, and preprocessing pipelines. All preprocessing steps were standardized across experimental conditions and classifier architectures. When processing full H&E images, we divided them into fixed-size patches and encoded each patch individually. During training, all patch embeddings were aggregated into a single global batch to avoid batch effects and ensure fair comparison with direct extrapolation. During inference, patch embeddings were processed per sample.

### 4.7. Implementation of HOPE

HOPE employs a two-layer linear projection with intermediate auxiliary supervision for the fusion model component, where auxiliary labels correspond to spatial omics-derived TME signatures. The fusion model computes: (1) an auxiliary projection from input H&E embeddings to spatial omics-derived signature logits, forming auxiliary features; and (2) a target projection from auxiliary features to prognosis logits, with both outputs normalized via softmax to yield respective probabilities.

Specifically, given input embeddings 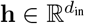, the auxiliary projection is defined as:

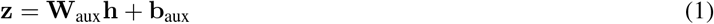

where 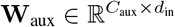 and 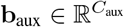, with *C*_aux_ denoting the number of spatial omics-derived signatures. The auxiliary features **z** are passed through softmax to obtain patch-level signature probabilities 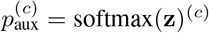.

The target projection follows as:

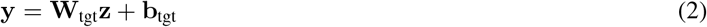

where 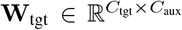and 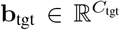, with *C*_tgt_ denoting the number of prognosis classes. The output **y** is similarly normalized via softmax to yield patient-level prognosis probabilities 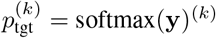.

The loss function integrates spatial omics-derived signature and prognosis losses with weight regularization. For a batch of *N* patches, let 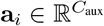 and 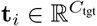 denote the ground-truth spatial omics-derived signature for patch *i* and the prognosis label of the corresponding patient, respectively. The spatial omics-derived signature loss is formulated as:

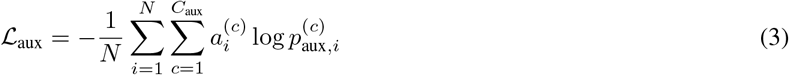

The prognosis loss is formulated as:

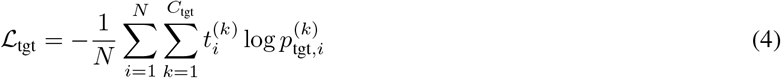

Weight regularization is applied with a dimension-scaled constant *c*_*𝓁*_:

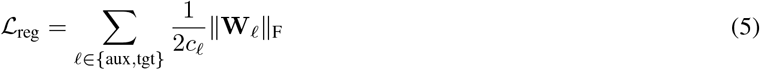

The total loss combines these terms:

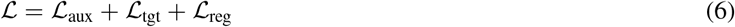

### 4.8. Implementation of direct extrapolation

Direct extrapolation serves as a baseline approach that omits the intermediate spatial omics-derived signature supervision present in HOPE. The model computes a direct projection from input H&E embeddings to prognosis class logits:

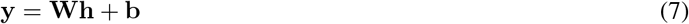

where 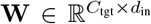 and 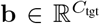, with *d*_in_ denoting the input embedding dimension and *C*_tgt_ the number of prognosis classes. The output **y** is normalized via softmax to obtain patient-level prognosis probabilities 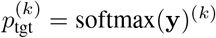.

The loss function integrates prognosis prediction cross-entropy with weight regularization. For a batch of *N* patches, let 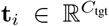 denote the ground-truth prognosis label for patch *i*. The prognosis loss follows Equation 4, and the weight regularization follows Equation 5 (applied to the single projection layer). The total loss combines these terms:

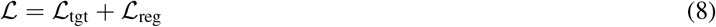

Both HOPE and direct extrapolation models operate at the patch level during training and inference. TME cluster predictions are computed independently per patch, whereas patient-level prognosis prediction is derived via median aggregation of patch-level probabilities across all H&E patches from each sample.

### 4.9. Cross validation and external validation

In the cross validation experiments of UPMC-HNC, to prevent data leakage across patients, we performed four-fold group cross validation with patient-level stratification. Within each fold, spatial omics signatures were derived exclusively from MIF data in the training split, while inference was restricted to H&E images in the held-out test split. To assess the generalizability of the HOPE framework, we conducted ablation experiments using H&E embeddings extracted from four foundation models: UNI2-h, MUSK, Phikon-v2, and PLIP.

HOPE model configurations were adapted for each feature extractor. For UNI2-h, MUSK, and Phikon-v2, batch normalization was incorporated following the auxiliary projection:

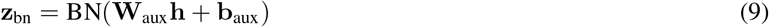

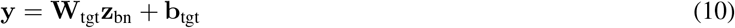

These variants were optimized using L-BFGS for one epoch at learning rate 1.0. For PLIP, a learnable temperature scaler was applied to the target projection output:

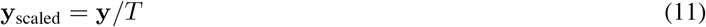

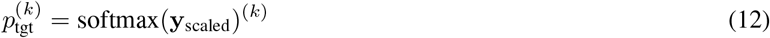

where *T* denotes a learned parameter. This configuration was first optimized with Adam for 5 epochs at learning rate 0.01, followed by L-BFGS for one epoch at learning rate 0.1.

For external validation on HANCOCK, all UPMC-HNC MIF and H&E data were utilized for spatial omics signature extraction and HOPE training. The resulting TME clusters were largely similar. We used UNI2-h to extract embeddings, with HOPE model configurations kept identical to those optimized during UPMC-HNC cross-validation. Independent testing was performed on the three predefined HANCOCK test sets (in-distribution, out-of-distribution, and oropharynx) [38] with three replicate experiments per set.

For the HCC immunotherapy response cohort, UNI2-h embeddings were extracted and HOPE was trained using Adam for 500 epochs at learning rate 0.01, with TME signatures annotated on the MIF data of the discovery cohort serving as auxiliary supervision. External validation on MDACC-HCC and UHT-HCC applied this trained model without modification.

All direct extrapolation baselines were optimized with L-BFGS for one epoch at learning rate 1.0, with identical data splits and following the same H&E image preprocessing, feature encoding, and inference protocols as HOPE.

### 4.10. Patient stratification by TME clusters

HOPE predicts patch-level TME clusters on H&E images. For stratification, each patch was assigned to the TME cluster with the highest predicted probability, and the proportion of patches belonging to each TME cluster was calculated per sample.

For the HNC cohorts, patients were stratified into high- and low-abundance groups based on the median proportion of the TME cluster of interest. Kaplan-Meier survival curves were generated for both groups. Among the HCC validation cohorts, overall survival data was available only for UHT-HCC and was included in survival analysis. Patches with maximum predicted probability ≤0.6 were excluded to ensure confident assignments. To benchmark TME cluster stratification against ground truth treatment response labels, we defined high/low abundance cutoffs based on the observed responder ratio: TME cluster proportions were ranked and partitioned to match the cohort’s responder-to-non-responder ratio. For example, in UHT-HCC (22 responders, 13 non-responders), My-T—a negative-response signature—was stratified into 13 high- and 22 low-abundance samples to match the non-responder-to-responder ratio (Fig. S7B).

## 5. Data and code availability

The HOPE source code is available at https://github.com/wangtyi/HOPE. The UPMC MIF and H&E datasets are available at https://zenodo.org/records/13179600 and https://zenodo.org/records/19163305, respectively. The HCC discovery dataset is available at https://zenodo.org/records/19123188. The HAN-COCK dataset is publicly available at https://hancock.research.fau.eu/download. The HCC validation dataset from MDACC and UHT are not publicly available due to institutional restrictions and data sharing agreements.

## Conflict of interest

M.B., A.T.M., A.E.T., and Z.W. are employees or former employees of Enable Medicine, Inc. C.M.S. is a cofounder, shareholder and employee of Vicinity Bio GmbH, and is a scientific advisor to and has received research funding from Enable Medicine Inc. All disclosed relationships are outside the scope of the submitted work.

## References

[1] Amelie Echle, Niklas Timon Rindtorff, Titus Josef Brinker, Tom Luedde, Alexander Thomas Pearson, and Jakob Nikolas Kather. Deep learning in cancer pathology: a new generation of clinical biomarkers. British journal of cancer, 124(4):686–696, 2021. 1

[2] Richard J Chen, Tong Ding, Ming Y Lu, Drew FK Williamson, Guillaume Jaume, Andrew H Song, Bowen Chen, Andrew Zhang, Daniel Shao, Muhammad Shaban, et al. Towards a general-purpose foundation model for computational pathology. Nature Medicine, 30(3):850–862, 2024. 1, 2, 5, 12

[3] Alexandre Filiot, Paul Jacob, Alice Mac Kain, and Charlie Saillard. Phikon-v2, a large and public feature extractor for biomarker prediction. arXiv, 2024. doi: 10.48550/arXiv.2409.09173. 1, 5, 12

[4] Jinxi Xiang, Xiyue Wang, Xiaoming Zhang, Yinghua Xi, Feyisope Eweje, Yijiang Chen, Yuchen Li, Colin Bergstrom, Matthew Gopaulchan, Ted Kim, et al. A vision–language foundation model for precision oncology. Nature, 638(8051):769–778, 2025. 1, 5, 12

[5] Zhi Huang, Federico Bianchi, Mert Yuksekgonul, Thomas J Montine, and James Zou. A visual–language foundation model for pathology image analysis using medical twitter. Nature medicine, 29(9):2307–2316, 2023. 1, 5, 12

[6] Andrew H Song, Guillaume Jaume, Drew FK Williamson, Ming Y Lu, Anurag Vaidya, Tiffany R Miller, and Faisal Mahmood. Artificial intelligence for digital and computational pathology. Nature Reviews Bioengineering, 1(12):930–949, 2023. 2

[7] Patrik L Stå hl Fredrik Salmén, Sanja Vickovic, Anna Lundmark, JoséFernández Navarro, Jens Magnusson, Stefania Giacomello, Michaela Asp, Jakub O Westholm, Mikael Huss, et al. Visualization and analysis of gene expression in tissue sections by spatial transcriptomics. Science, 353(6294): 78–82, 2016. 2

[8] Lambda Moses and Lior Pachter. Museum of spatial transcriptomics. Nature methods, 19(5):534–546, 2022. 2

[9] Yury Goltsev, Nikolay Samusik, Julia Kennedy-Darling, Salil Bhate, Matthew Hale, Gustavo Vazquez, Sarah Black, and Garry P Nolan. Deep profiling of mouse splenic architecture with codex multiplexed imaging. Cell, 174(4):968–981, 2018. 2

[10] Emma Lundberg and Georg HH Borner. Spatial proteomics: a powerful discovery tool for cell biology. Nature Reviews Molecular Cell Biology, 20 (5):285–302, 2019. 2

[11] Longqi Liu, Ao Chen, Yuxiang Li, Jan Mulder, Holger Heyn, and Xun Xu. Spatiotemporal omics for biology and medicine. Cell, 187(17):4488–4519, 2024. 2

[12] Aiwei Lin Ji, Adam J Rubin, Kristine Thrane, Siyuan Jiang, Daniel L Reynolds, Ryan M Meyers, Michael G Guo, Braden M George, Annelie Mollbrink, et al. Multimodal analysis of composition and spatial architecture in human squamous cell carcinoma. Cell, 182(2):497–514.e22, 2020. 2

[13] Karin Pelka, Matan Hofree, Jonathan H Chen, Siranush Sarkizova, John D Pirl, Vjola Jorgji, Arman Bejnood, Danielle Dionne, Wenqi Ge, Kristen H Xu, et al. Spatially organized multicellular immune hubs in human colorectal cancer. Cell, 184(18):4734–4752.e20, 2021.

[14] Sabrina M Lewis, Marie-Liesse Asselin-Labat, Quan Nguyen, Jean Berthelet, Xiao Tan, Verena C Wimmer, Delphine Merino, Kelly L Rogers, and Shalin H Naik. Spatial omics and multiplexed imaging to explore cancer biology. Nature methods, 18(9):997–1012, 2021.

[15] Christian M Schürch, Salil S Bhate, Graham L Barlow, Darci J Phillips, Luca Noti, Inti Zlobec, Pauline Chu, Sarah Black, Janos Demeter, David R McIlwain, et al. Coordinated cellular neighborhoods orchestrate antitumoral immunity at the colorectal cancer invasive front. Cell, 182(5):1341–1359, 2020. 2

[16] Jia-Ren Lin, Yu-An Chen, Daniel Campton, Jeremy Cooper, Shannon Coy, Clarence Yapp, Juliann B Tefft, Erin McCarty, Keith L Ligon, Scott J Rodig, et al. High-plex immunofluorescence imaging and traditional histology of the same tissue section for discovering image-based biomarkers. Nature cancer, 4(7):1036–1052, 2023. 2

[17] Xiao Qian Wang, Esther Danenberg, Chiun-Sheng Huang, Daniel Egle, Maurizio Callari, Begoña Bermejo, Matteo Dugo, Claudio Zamagni, Marc Thill, Anton Anton, et al. Spatial predictors of immunotherapy response in triple-negative breast cancer. Nature, 621(7980):868–876, 2023.

[18] Thazin N Aung, James Monkman, Jonathan Warrell, Ioannis Vathiotis, Katherine M Bates, Niki Gavrielatou, Ioannis P Trontzas, Chin Wee Tan, Aileen I Fernandez, Myrto Moutafi, et al. Spatial signatures for predicting immunotherapy outcomes using multi-omics in non-small cell lung cancer. Nature Genetics, 57(10):2482–2493, 2025.

[19] Zhenqin Wu, Alexandro E Trevino, Eric Wu, Kyle Swanson, Honesty J Kim, H Blaize D’Angio, Ryan Preska, Gregory W Charville, Piero D Dalerba, Ann Marie Egloff, et al. Graph deep learning for the characterization of tumour microenvironments from spatial protein profiles in tissue specimens. Nature Biomedical Engineering, 6(12):1435–1448, 2022. 2, 12

[20] Zhenqin Wu, Joseph Boen, Sonali Jindal, Sreyashi Basu, Matthew Bieniosek, Siyu He, Michael LaPelusa, Aaron T Mayer, Ahmed O Kaseb, James Zou, et al. Spatial multi-omics and deep learning reveal fingerprints of immunotherapy response and resistance in hepatocellular carcinoma. bioRxiv, 2025. 2, 8, 11, 12

[21] Jian Hu, Xiangjie Li, Kyle Coleman, Amelia Schroeder, Nan Ma, David J Irwin, Edward B Lee, Russell T Shinohara, and Mingyao Li. Spagcn: Integrating gene expression, spatial location and histology to identify spatial domains and spatially variable genes by graph convolutional network. Nature methods, 18(11):1342–1351, 2021. 2

[22] Weiqing Chen, Pengzhi Zhang, Tu N Tran, Yiwei Xiao, Shengyu Li, Vrutant V Shah, Hao Cheng, Kristopher W Brannan, Keith Youker, Li Lai, et al. A visual–omics foundation model to bridge histopathology with spatial transcriptomics. Nature Methods, 22(7):1568–1582, 2025. 2

[23] Kyle Coleman, Amelia Schroeder, Melanie Loth, Daiwei Zhang, Jeong Hwan Park, Ji-Youn Sung, Niklas Blank, Alexis J Cowan, Xuyu Qian, Jianfeng Chen, et al. Resolving tissue complexity by multimodal spatial omics modeling with miso. Nature methods, 22(3):530–538, 2025. 2

[24] Yuansong Zeng, Zhuoyi Wei, Weijiang Yu, Rui Yin, Yuchen Yuan, Bingling Li, Zhonghui Tang, Yutong Lu, and Yuedong Yang. Spatial transcriptomics prediction from histology jointly through transformer and graph neural networks. Briefings in Bioinformatics, 23(5):bbac297, 2022. 2

[25] Xiaohang Fu, Yue Cao, Beilei Bian, Chuhan Wang, Dinny Graham, Nirmala Pathmanathan, Ellis Patrick, Jinman Kim, and Jean Yee Hwa Yang. Spatial gene expression at single-cell resolution from histology using deep learning with ghist. Nature methods, 22(9):1900–1910, 2025.

[26] Chuhan Wang, Adam S Chan, Xiaohang Fu, Shila Ghazanfar, Jinman Kim, Ellis Patrick, and Jean YH Yang. Benchmarking the translational potential of spatial gene expression prediction from histology. Nature Communications, 16(1):1544, 2025. 2

[27] Zhe Li, Yuchen Li, Jinxi Xiang, Xiyue Wang, Sen Yang, Xiaoming Zhang, Feyisope Eweje, Yijiang Chen, Xiangde Luo, Yuanyuan Li, et al. Ai-enabled virtual spatial proteomics from histopathology for interpretable biomarker discovery in lung cancer. Nature Medicine, 32:231–244, 2026. 2

[28] Jeya Maria Jose Valanarasu, Hanwen Xu, Naoto Usuyama, Chanwoo Kim, Cliff Wong, Peniel Argaw, Racheli Ben Shimol, Angela Crabtree, Kevin Matlock, Alexandra Q Bartlett, et al. Multimodal ai generates virtual population for tumor microenvironment modeling. Cell, 189(2):386–400, 2026. 2

[29] Peng Zhang, Chaofei Gao, Zhuoyu Zhang, Zhiyuan Yuan, Qian Zhang, Ping Zhang, Shiyu Du, Weixun Zhou, Yan Li, and Shao Li. Systematic inference of super-resolution cell spatial profiles from histology images. Nature Communications, 16(1):1838, 2025. 2

[30] Zhe Li, Seyed Hossein Mirjahanmardi, Rasoul Sali, Feyisope Eweje, Matthew Gopaulchan, Leon Kloker, Xiaoming Zhang, Guoxin Li, Yuming Jiang, and Ruijiang Li. Automated cell annotation and classification on histopathology for spatial biomarker discovery. Nature Communications, 16(1):6240, 2025.

[31] Xiao Tan, Onkar Mulay, Jacky Xie, Samual MacDonald, Taehyun Kim, Chenhao Zhou, Zherui Xiong, Samuel X Tan, Nan Ye, Amy McCart Reed, et al. Robust and interpretable prediction of gene markers and cell types from spatial transcriptomics data. Nature Communications, 17:1781, 2026. 2

[32] Muhammad Dawood, Kim Branson, Sabine Tejpar, Nasir Rajpoot, and Fayyaz ul Amir Afsar Minhas. Confounding factors and biases abound when predicting molecular biomarkers from histological images. Nature Biomedical Engineering, pages 1–15, 2026. 2

[33] Lichun Ma, Barbara Xiong, Meng Liu, and Kai Tan. Cellular neighborhoods in cancer. Nature Cancer, pages 1–13, 2026. 2

[34] Roy Noy and Jeffrey W Pollard. Tumor-associated macrophages: from mechanisms to therapy. Immunity, 41(1):49–61, 2014. 5

[35] Lei Chen, Shu-Cheng Wan, Liang Mao, Cong-Fa Huang, Lin-Lin Bu, and Zhi-Jun Sun. Nlrp3 in tumor-associated macrophages predicts a poor prognosis and promotes tumor growth in head and neck squamous cell carcinoma. Cancer Immunology, Immunotherapy, 72(6):1647–1660, 2023. 5

[36] Wolf Herman Fridman, Franck Pagés, Catherine Sautés-Fridman, and Jèrôme Galon. The immune contexture in human tumours: impact on clinical outcome. Nature Reviews Cancer, 12(4):298–306, 2012. 5

[37] Jean-Luc Teillaud, Ana Houel, Marylou Panouillot, Clèmence Riffard, and Marie-Caroline Dieu-Nosjean. Tertiary lymphoid structures in anticancer immunity. Nature Reviews Cancer, 24(9):629–646, 2024. 5

[38] Marion Dörrich, Matthias Balk, Tatjana Heusinger, Sandra Beyer, Hamed Mirbagheri, David J Fischer, Hassan Kanso, Christian Matek, Arndt Hartmann, Heinrich Iro, et al. A multimodal dataset for precision oncology in head and neck cancer. Nature Communications, 16(1):7163, 2025. 8, 14

[39] Ira Mellman, George Coukos, and Glenn Dranoff. Cancer immunotherapy comes of age. Nature, 480(7378):480–489, 2011. 8

[40] Tarana Gupta and Nikhil Sai Jarpula. Hepatocellular carcinoma immune microenvironment and check point inhibitors-current status. World Journal of Hepatology, 16(3):353, 2024. 8

[41] C Luchini, F Bibeau, MJL Ligtenberg, Navdeep Singh, A Nottegar, T Bosse, R Miller, N Riaz, J-Y Douillard, F Andre, et al. Esmo recommendations on microsatellite instability testing for immunotherapy in cancer, and its relationship with pd-1/pd-l1 expression and tumour mutational burden: a systematic review-based approach. Annals of Oncology, 30(8):1232–1243, 2019. 8

[42] Ahmed Omar Kaseb, Elshad Hasanov, Hop Sanderson Tran Cao, Lianchun Xiao, Jean-Nicolas Vauthey, Sunyoung S Lee, Betul Gok Yavuz, Yehia I Mohamed, Aliya Qayyum, Sonali Jindal, et al. Perioperative nivolumab monotherapy versus nivolumab plus ipilimumab in resectable hepatocellular carcinoma: a randomised, open-label, phase 2 trial. The lancet Gastroenterology & hepatology, 7(3):208–218, 2022. 8, 11

[43] Deniz Seyhan, Manon Allaire, Yaojie Fu, Filomena Conti, Xin Wei Wang, Bin Gao, and Fouad Lafdil. Immune microenvironment in hepatocellular carcinoma: from pathogenesis to immunotherapy. Cellular & Molecular Immunology, 22(10):1132–1158, 2025. 8

[44] Hongyu Zheng, Xueqiang Peng, Shuo Yang, Xinyu Li, Mingyao Huang, Shibo Wei, Sheng Zhang, Guangpeng He, Jiaxing Liu, Qing Fan, et al. Targeting tumor-associated macrophages in hepatocellular carcinoma: biology, strategy, and immunotherapy. Cell Death Discovery, 9(1):65, 2023. 8

[45] Fen Liu, Xianying Li, Yiming Zhang, Shan Ge, Zhan Shi, Qingbin Liu, and Shulong Jiang. Targeting tumor-associated macrophages to overcome immune checkpoint inhibitor resistance in hepatocellular carcinoma. Journal of Experimental & Clinical Cancer Research, 44(1):227, 2025. 8

[46] Abigail H Cleveland and Yi Fan. Reprogramming endothelial cells to empower cancer immunotherapy. Trends in molecular medicine, 30(2):126–135, 2024. 8

