## Supplementary Figures for "HOPE: Interpretable Histology Analysis with Spatial Omics-Derived Signatures for Precision Oncology"

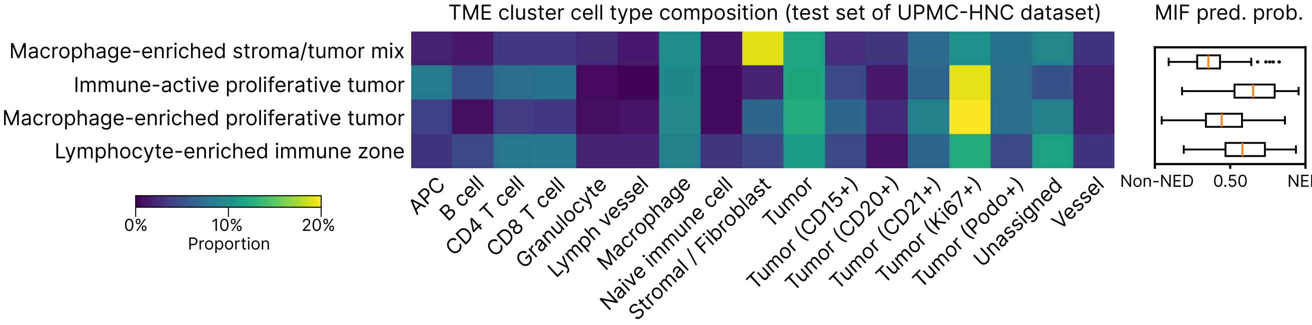


Fig. S1. **TME cluster cell type composition predicted by HOPE on the UPMC-HNC cross-validation test set.** Consistent with Fig. 2C, TME clusters were assigned to patches via H&E-only prediction, then mapped to spatially aligned MIF-based cell subgraphs. Cell type composition for each TME cluster was obtained by averaging cell type proportions across constituent cell subgraphs, without using SP features. The resulting composition closely matches that of the training set (Fig. 2C), preserving characteristic patterns and validating HOPE's reliability in TME cluster prediction. The right panel shows survival prediction probabilities from the trained SP model applied to these MIF-based cell subgraphs, with trends matching Fig. 2B.

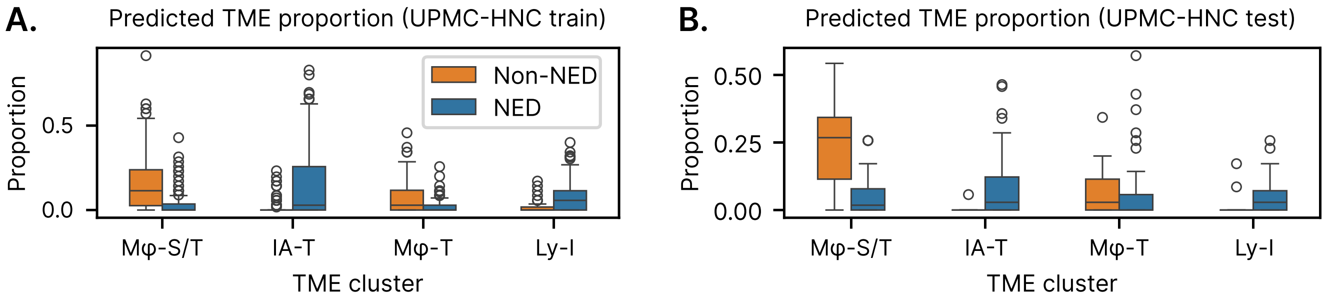


Fig. S2. **Proportion distributions of TME clusters across H&E patches per patient for UPMC-HNC cross validation. A.** The TME proportions in the training set, obtained by uniformly sampling patches via a sliding window using the trained HOPE model, are distinct from random sampling of ground truth regions. **B.** The predicted TME proportions on the test set from the same fold using the identical trained model. M$\varphi$-S/T and M$\varphi$-T were more prevalent in patients with negative outcomes (Non-NED), whereas IA-T and Ly-I were markedly more abundant in patients with positive outcomes (NED). Abbreviations: M$\varphi$-S/T, macrophage-enriched stroma/tumor mix; IA-T, immune-active proliferative tumor; M$\varphi$-T, macrophage-enriched proliferative tumor; Ly-I, lymphocyte-enriched immune zone.

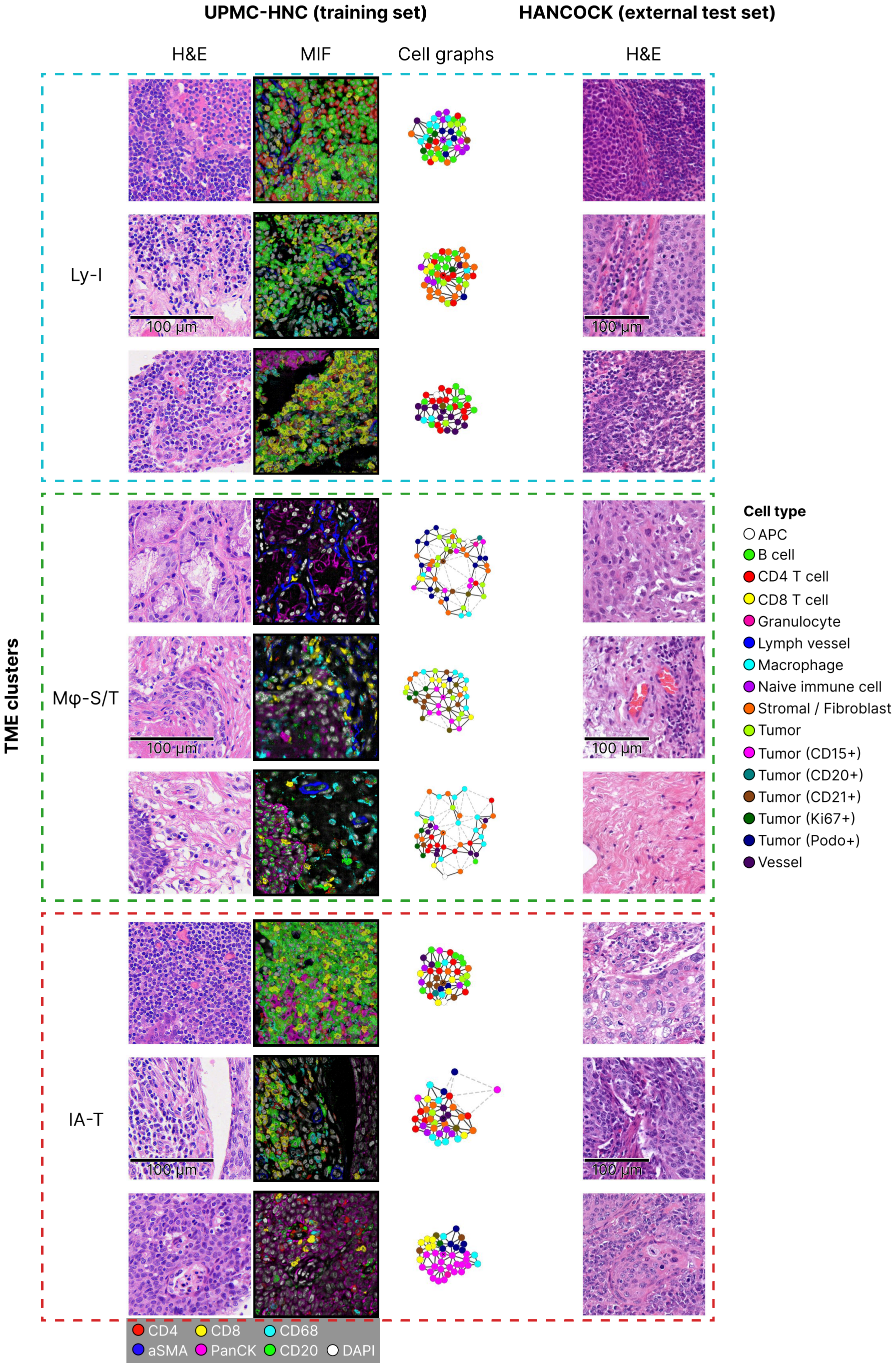


Fig. S3. **Examples of H&E, MIF patches, and cell graphs identified as Ly-I, M$\varphi$-S/T, and IA-T for HNC.** UPMC-HNC shows MIF images and cell graphs from which signatures were derived as ground truth, alongside spatially aligned H&E images that were used during training. HANCOCK demonstrates examples predicted by HOPE using H&E images alone. Abbreviations: Ly-I, lymphocyte-enriched immune zone; M$\varphi$-S/T, macrophage-enriched stroma/tumor mix; IA-T, immune-active proliferative tumor.

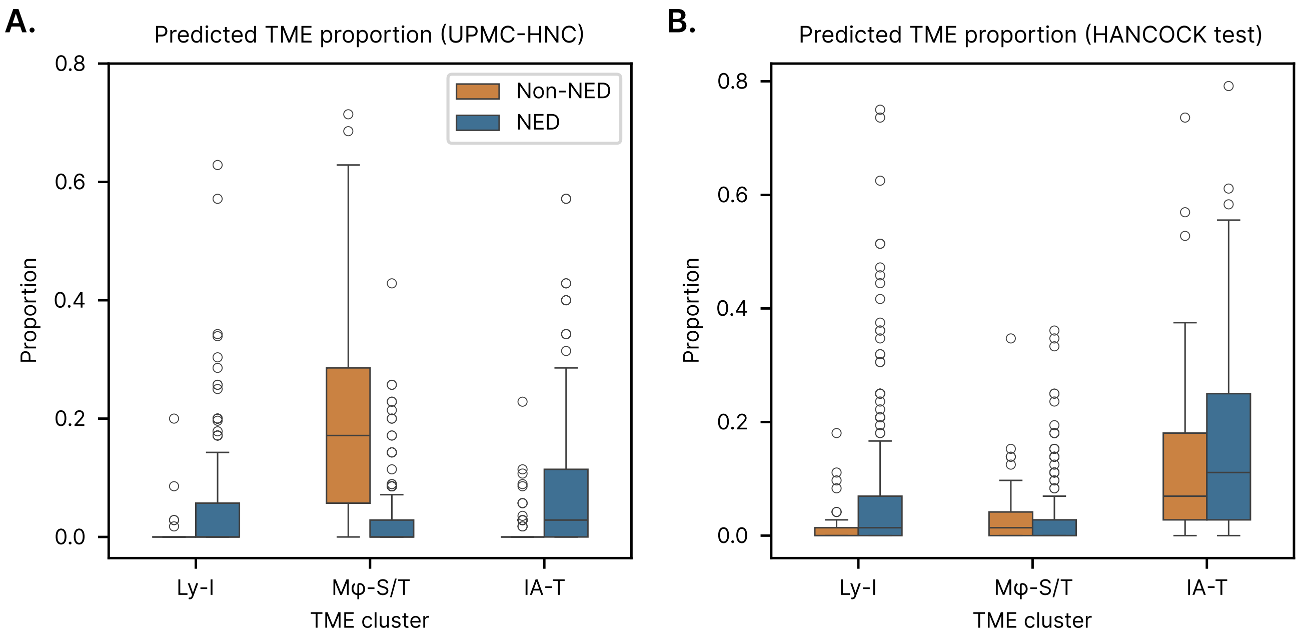


Fig. S4. **Proportion distributions of TME clusters across H&E patches per patient for HNC external validation. A.** The ground truth TME proportions from UPMC-HNC (training set). **B.** The predicted TME proportions for HANCOCK (test set) using only H&E. M$\varphi$-S/T was more prevalent in patients with negative outcomes (Non-NED), whereas Ly-I and IA-T were more prevalent in patients with positive outcomes (NED). Consistent patterns were observed in both UPMC-HNC and HANCOCK. Abbreviations: Ly-I, lymphocyte-enriched immune zone; M$\varphi$-S/T, macrophage-enriched stroma/tumor mix; IA-T, immune-active proliferative tumor.

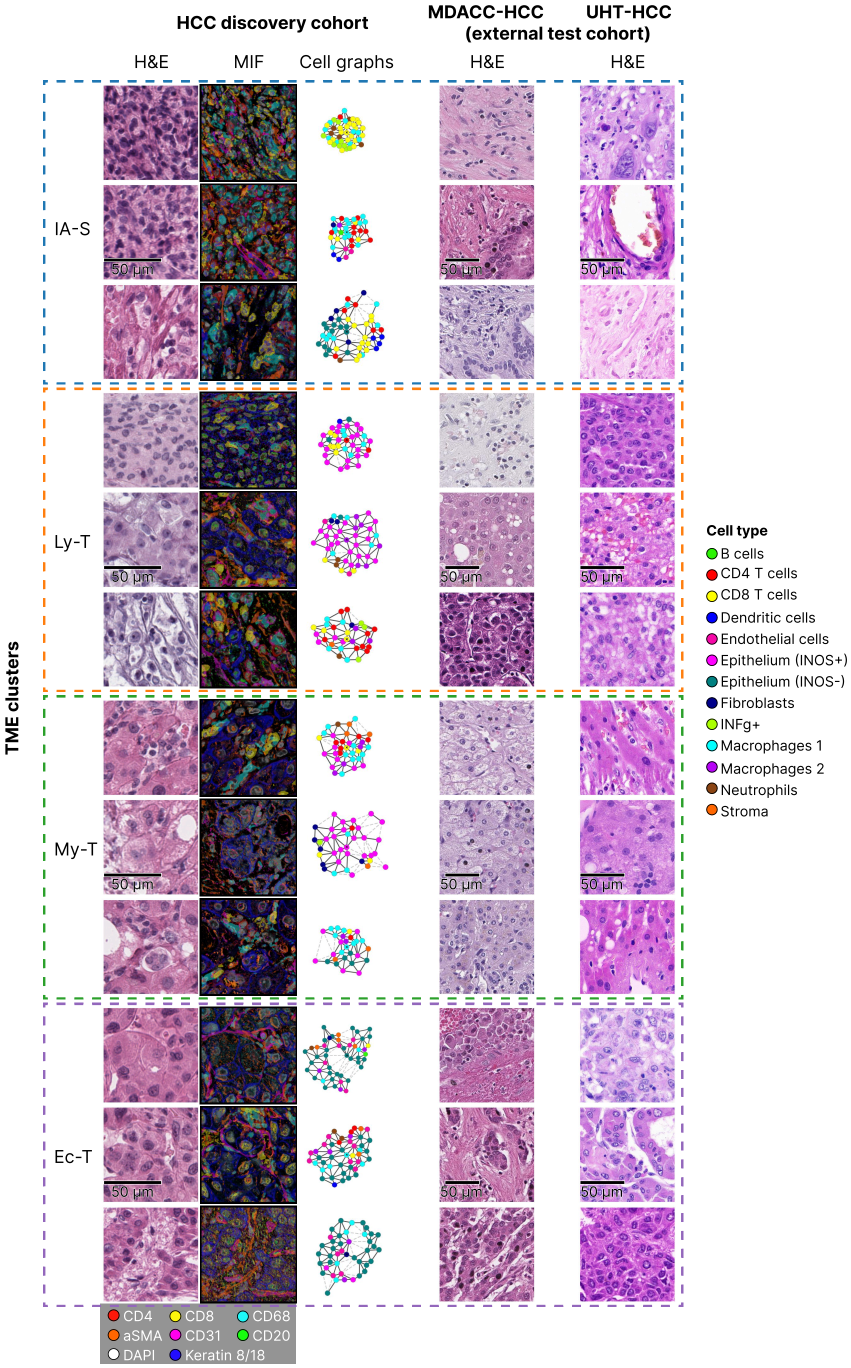


Fig. S5. **Examples of H&E, MIF patches, and cell graphs identified as IA-S, Ly-T, My-T, and Ec-T for HCC discovery and external test cohorts.** The HCC discovery cohort shows MIF images and cell graphs from which signatures were derived as ground truth, alongside spatially aligned H&E images. MDACC-HCC and UHT-HCC demonstrate examples from two external test cohorts predicted by HOPE using H&E images alone. Abbreviations: IA-S, immune-active stroma; Ly-T, lymphocyte-rich tumor; My-T, myeloid cell-rich tumor; Ec-T, endothelial cell-rich tumor.

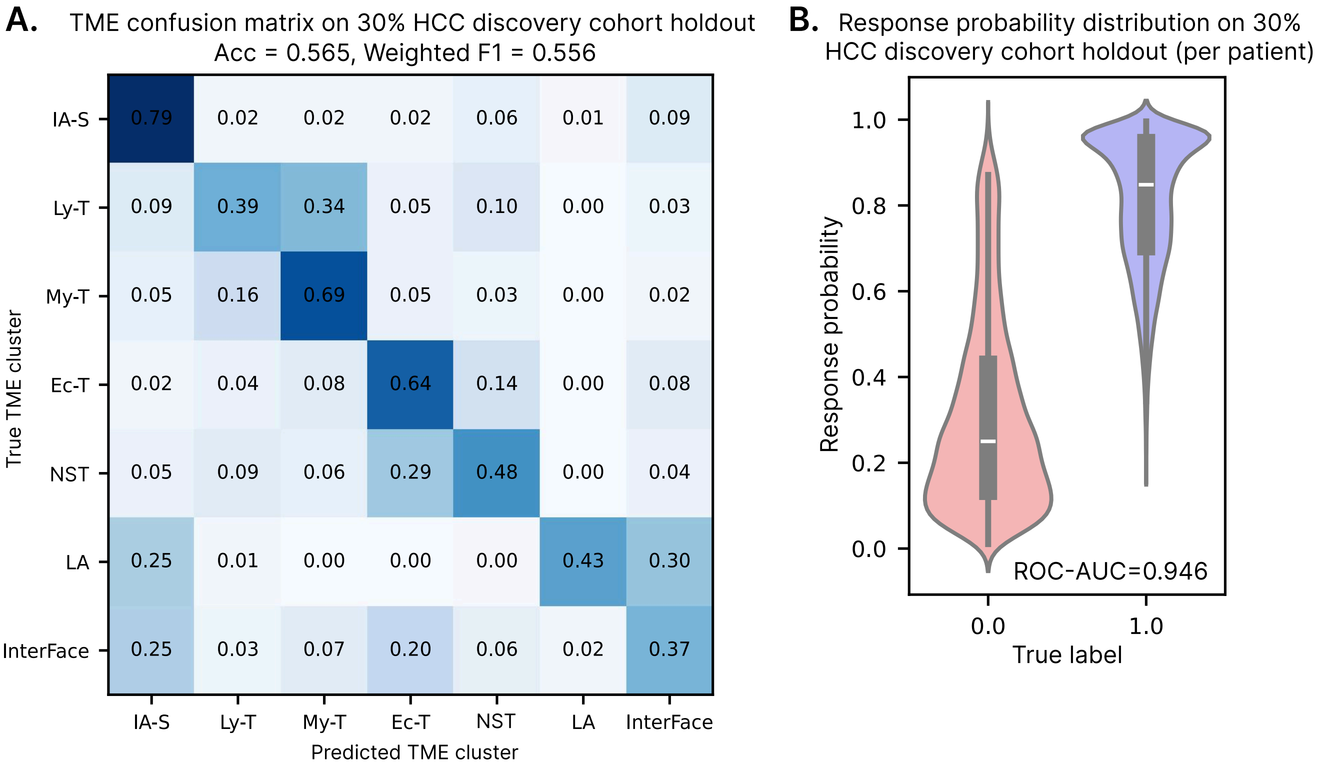


Fig. S6. **Internal validation on the HCC discovery cohort.** The HCC discovery cohort was split 7:3 into train and test sets for internal validation. **A.** TME clusters predicted by HOPE using H&E on the 30% discovery cohort holdout. Abbreviations: IA-S, immune-active stroma; Ly-T, lymphocyte-rich tumor; My-T, myeloid cell-rich tumor; Ec-T, endothelial cell-rich tumor; NST, non-specific tumor; LA, lymphoid aggregates; InterFace, LA-tumor interface. **B.** Distribution of immunotherapy response probabilities predicted by HOPE using H&E for patients in the 30% discovery cohort holdout.

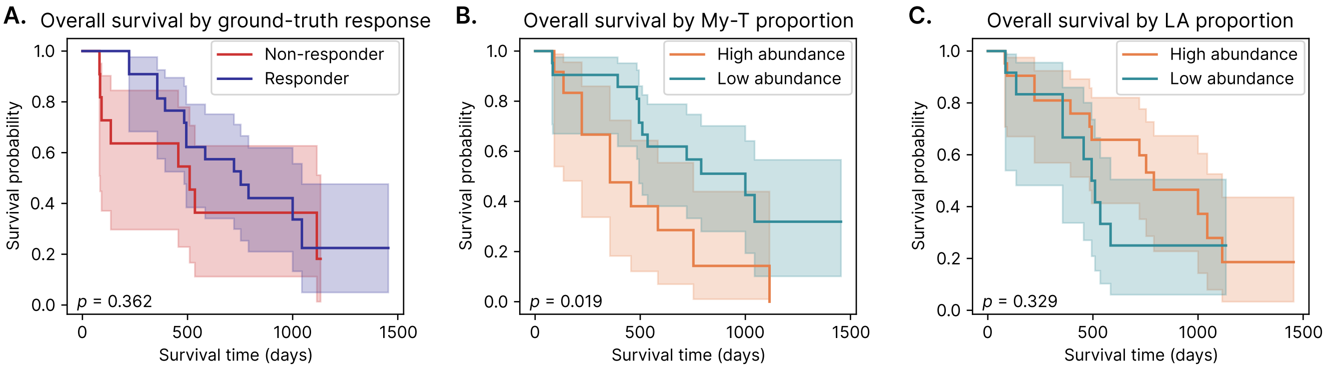


Fig. S7. **KM stratification of patients in the UHT-HCC dataset. A.** Stratification by ground-truth immunotherapy response labels (22 Responder, 13 Non-responder). **B.** Stratification by predicted My-T (myeloid cell-rich tumor) abundance (22 Low abundance, 13 High abundance). **C.** Stratification by predicted LA (lymphoid aggregates) abundance (13 Low abundance, 22 High abundance). All predictions were based on H&E images without omics data. Equal-sized high- and low-risk groups across all three panels enable direct comparison of prognostic stratification performance.
